# Covalent Inhibition of New Delhi Metallo-β-Lactamases NDM-1 and NDM-5 by 3-Bromopyruvate

**DOI:** 10.64898/2026.06.10.731408

**Authors:** Jack K. Bradley, Karina Calvopina Tapia, Sabrina J. Moyo, Ellinor Shore, Peter Nambala, W. David Hong, Christopher J. Schofield, Adam P. Roberts

**Author notes:** Corresponding Author: Adam Roberts,.

## Abstract

Resistance to β-lactam antibiotics, including carbapenems, mediated by metallo-β-lactamases (MBLs), including the New Delhi metallo-β-lactamase (NDM) MBL subfamily, is increasing. No MBL inhibitors are currently approved for clinical use with most reported MBL inhibitors are metal ion chelators, acting either at the Zn(II) ion active site and/or in solution. The hexokinase inhibitor 3-bromopyruvate (3-BP) is reported to inhibit NDM-1. We found that 3-BP selectively restored the antimicrobial activity of meropenem against carbapenem resistant *Escherichia coli, Klebsiella pneumoniae* and *Acinetobacter baumannii* strains, obtained from clinical and environmental isolates from Tanzania and Malawi, containing genes that encode NDM-1 or NDM-5, but not against strains containing genes encoding for serine β-lactamases. Mass spectrometry studies with NDM-1 and NDM-5 support a mechanism involving covalent reaction of 3-BP with an active site cysteine residue. The results will promote work on the development of covalently reacting MBL inhibitors, a strategy that has been successful for inhibition of the nucleophilic serine β-lactamases.

## Introduction

Antimicrobial resistance (AMR) is an increasing global health problem, with an estimated 4.95 million AMR associated deaths in 2019.^1, 2^ Resistance to carbapenems, once reserved as a last resort treatment against multi-drug resistant bacteria is now identified as a critical priority by the World Health Organisation. The β-lactams, including penicillins, cephalosporins and carbapenems, are the most widely prescribed antibiotic class. The targets of β-lactams are transpeptidases (penicillin-binding proteins, PBPs), responsible for the crosslinking of peptidoglycan during bacterial cell wall biosynthesis.^3^ Inhibition of PBPs prevents the protein from performing its normal function, blocking crosslinking and resulting in weakening of the cell wall and autolysis.^4, 5^ An important mechanism of resistance to β-lactams involves their β-lactamase catalysed hydrolysis, rendering the antibiotic inactive.^6, 7^ Ambler class A, C and D β-lactamases are nucleophilic serine enzymes, with the class B metallo β-lactamases (MBLs) requiring one or two active site Zn^2+^ ions that activate a water molecule for β-lactam hydrolysis. MBLs are divided into subclasses B1–B3 based on the number of metal ions found in the active site as well sequence and structural similarities, with the B1 MBL subfamily currently being the most clinically relevant. ^8, 9^

Following the discovery of the subclass B1 MBL New Delhi metallo-β-lactamase-1 (NDM-1), NDM-1 and its multiple variants have emerged as one of the most prominent and clinically relevant MBL subfamilies.^10, 11^ They are found to confer resistance to nearly all β-lactams, including carbapenems.^12-14^ NDMs are now prevalent in Asia, the Middle East, Europe, Africa and the Americas, largely driven by mobile plasmids that transfer between Gram-negative bacteria such as *Escherichia coli, Klebsiella pneumoniae* and *Acinetobacter baumannii*.^15^ The active site of NDM-1 employs two Zn(II) ions for catalysis. In the NDM zinc-1 site, the zinc ion is coordinated by three histidine-residues, whilst at the zinc-2 site, the zinc ion is coordinated to by a cysteine, a histidine and an aspartic acid residue.^16^ The cysteine residue is conserved across B1 MBL active sites and may possess sufficient nucleophilicity to enable covalent targeting.

Clinically, serine β-lactamase inhibitors (e.g. clavulanic acid) work by covalently reacting mechanisms and are used in combination with β-lactam antibiotics to overcome resistance, by contrast, no clinically useful MBL inhibitors are currently available.^17, 18^ MBLs are challenging targets due to their structural diversity and the presence of structurally related human MBL fold enzymes.^10, 19^ Most reported MBL inhibitors are metal ion chelators, acting either at the Zn(II) ion active site and/or in solution, with the known hexokinase inhibitor 3-bromopyruvate (3-BP) being an exception.^20, 21^ 3-Bromopyruvate (3-BP) is an small electrophilic halomethyl ketone derivative of the cellular metabolite pyruvate and has previously been explored as a potential anticancer treatment.^22^ 3-BP has demonstrated activity in a number of animal tumour models and has been used in clinical trials.^23, 24^ These positive results, combined with low cytotoxicity versus mouse fibroblastic cells at concentrations up to 200 μM, suggests 3-BP is a potentially safe lead for therapeutic development.

3-BP is also reported to inhibit NDM-1, restoring the activities of meropenem and cefazolin against *E. coli* strains producing NDM-1 (at 64 mg/L).^21^ 3-BP was shown to inhibit subclass B1 MBLs (NDM-1, VIM-2, IMP-1) and B2 (ImiS) MBLs, but not the B3 MBL L1, in accord with presence of an active site cysteine residue in the B1 and B2 MBLs, but not the B3 MBLs.^21,25^

Here we report that moderate concentrations of 3-BP restore the activity of meropenem at the susceptibility breakpoint (2 mg/L) against clinically and environmentally derived resistant Enterobacterales isolates containing MBLs NDM-1 and NDM-5. 3-BP, however, did not restore activity of amoxicillin at the susceptibility breakpoint (8 mg/L) against environmentally derived Enterobacterales isolates containing SBLs TEM-1B, CTX-M-15, ACT-16, OXA-1 or OXA-48. The parent scaffold pyruvic acid showed no activity against isolates containing MBLs or SBLs, with the introduction of bromine in 3-BP leading to the observed inhibition. Small molecule mass spectrometry (MS) studies, followed by denaturing mass spectrometry studies on apo- and metallated-forms of NDM-1 and NDM-5 to investigate the mechanisms of inhibition by 3-BP, identifying the formation of covalent adducts.

## Materials and methods

Mueller Hinton broth, amoxicillin and meropenem were from Fisher Scientific. All AST methods follow European Committee on Antimicrobial Susceptibility Testing (EUCAST) guidelines. Bacterial strains were stored in glycerol stocks at -70 °C. 3-Bromopyruvate and pyruvic acid (> 98 % purity) were from Sigma Aldrich. Cysteine and glutathione (> 95 % purity) were from Fluorochem.

### Broth microdilution assays

Selected bacterial strains were grown from glycerol stocks on Mueller Hinton agar for 16-18 hours overnight at 37°C. Selected colonies were then suspended in 10 mL Mueller Hinton broth and allowed to grow for 16-18 hours overnight at 37°C. The optical density (OD) of the inoculum was measured and then the inoculum diluted to give an optical density of 1 (OD1) at a 600 nm wavelength. OD1 was then further diluted 1/1000 in Mueller Hinton broth to obtain an inoculum suspension equivalent to for a single drug MIC experiment or 1/500 inoculum suspension equivalent for a rescue concentration experiment. Once prepared the inoculum was used within 30 minutes. Serial dilution was performed from wells 2-10 to obtain final inhibitor concentrations from 256 mg/L to 1 mg/L with well 11 acting as a control (no drug). To determine the rescue concentration of 3-BP and pyruvic acid, meropenem or amoxicillin was then added to each well at a final concentration of 2 mg/L or 8 mg/L respectively when combined with 3-BP or pyruvic acid. After inoculation, the 96-well plates were incubated at 37°C for 16–18 hours. All drugs were tested via three biological replicates and three technical replicates performed. MIC/RC was recorded at the lowest concentration of no visible growth of bacteria.

### Mass spectrometry assays

#### Small molecule mass spectrometry studies

For small molecule mass spectrometry studies 10mM stock concentrations of 3-BP, glutathione and cysteine were made using HPLC grade water. 2 ml of 3-BP was added to 2 ml volumes of glutathione and cysteine and allowed to stir for 18 hours at 37 °C. After 18 hours the reaction mixture was filtered and submitted for mass spectrum analysis. High resolution mass spectra (HRMS) were obtained using an Agilent QTOF 6500 series spectrometer, equipped with an electrospray ionisation interface and operated in the positive mode.

#### Preparation of the metal-depleted enzyme

The zinc content of the stock enzyme preparation was depleted by dialysis in a MINI dialysis device (0.5 mL, 10 kDa MWCO) against a buffer containing 50 mM HEPES (pH 7.2), 30 mM EDTA, and 5 mM phenanthroline. MBL protein was subsequently dialysed against 50 mM HEPES (pH 7.2), 1% Chelex resin to remove free metal ions metals and phenanthroline. Following dialysis, the protein was concentrated using an Amicon® Ultra Centrifugal Filter (0.5 Ml, 10 kDa MWCO)

#### ESI-mass spectrometry

Purified NDM-1/NDM-5 and metal depleted NDM-1/NDM-5 stocks were diluted to 1 μM and 10 equivalents of 3-BP was added to each sample. Samples were analysed by LC–ESI mass spectrometry using a Xevo G2-S mass spectrometer coupled to an Acquity UPLC system (Waters) fitted with a ProSwift RP-4H column (1 × 50 mm; Thermo Fisher Scientific). Proteins were loaded in 95% water, 5% acetonitrile, and 0.1% formic acid and eluted over a 10 min gradient to 95% acetonitrile containing 0.1% formic acid before direct introduction into the ESI source. Protein retention times were approximately 4–5 min. Data were processed using MassLynx 4.1 (Waters), and spectra were deconvoluted using the MaxEnt1 algorithm.

#### Protein modelling

AlphaFold server (version 3) was used for automated protein tertiary structure modelling of apo-NDM-1 and NDM-5.^26^ Modelling results were visualised using UCSF Chimera X 1.10.1.^27^

## Results and discussion

### Minimum inhibitory concentration & rescue concentration

To investigate if 3-BP restored antimicrobial activity of β-lactam antibiotics against bacterial strains producing NDM-1 and NDM-5, minimum inhibitory concentrations (MICs) of meropenem, pyruvic acid and 3-BP were determined (**Table 1A**) according to the European Committee on Antimicrobial Susceptibility Testing (EUCAST) guidelines.^28^ Clinical and environmental isolates of *A. baumannii, K. pneumoniae* and *E. coli* producing NDM-1 or NDM-5 were obtained during studies investigating neonatal diseases in Tanzania.^15, 29, 30^ Additional isolates were obtained through the National Collection of Type Cultures and noted by their NCTC number or from an ongoing hospital wastewater surveillance study (STRESST) based at the Queen Elizabeth Central Hospital, Blantyre, Malawi. It is notable that genes coding for serine β-lactamases are also in the different tested isolates, including *bla*_ADC_, *bla*_CARB_, *bla*_CTX_, *bla*_TEM_ and *bla*_OXA_ (**SI-1**).^15, 29^

**Table 1.** (**A**) Microbiological comparison of meropenem, pyruvic acid and 3-bromopyruvate activities against NDM-1 or NDM-5 containing *A. baumannii, K. pneumoniae and E. coli* strains. MIC: minimum concentration required for no visible growth of bacteria. (**B**) Concentrations of pyruvic acid and 3-BP needed to restore the activity of meropenem at its EUCAST susceptibility breakpoint concentration of 2 mg/L in NDM-1 or NDM-5 containing *A. baumannii, K. pneumoniae* and *E. coli* strains. RC: minimum concentration required for no visible growth of bacteria when used synergistically with meropenem set at a concentration of 2 mg/L. Plates were incubated at 37°C for 16–18 h. Three biological and three technical replicates were performed.*-Table 1B, value of > 256 mg/L corresponds to no observable recovery of meropenem activity at 2 mg/L.

| A - MIC (mg/L) |  |  |  |  |  |  |  |  |  |  |  |
| --- | --- | --- | --- | --- | --- | --- | --- | --- | --- | --- | --- |
|  | *Acinetobacter baumannii* |  |  | *klebsiella pneumoniae* |  |  |  | *Escherichia coli* |  |  |  |
| Compound | NCTC 12156 (No MBL) | DT05 44 (NDM -1) | DT011 39 (NDM-1) | NCTC134 42 (No MBL) | NTC134 43 (NDM-1) | ArmMLGA 211 (NDM-1) | ESSCAI1 22 (NDM-1) | SM247682 033 (No MBL) | MLGA 91 (NDM-5) | MLGA 92 (NDM-5) | ESCAI2 31 (NDM-5) |
| Meropenem | < 1 | 128 | 128 | < 1 | > 256 | 128 | 128 | < 1 | 128 | 128 | 64 |
| Pyruvic acid | > 256 | > 256 | > 256 | > 256 | > 256 | > 256 | > 256 | > 256 | > 256 | > 256 | > 256 |
| 3-Bromopyruvate | > 256 | > 256 | > 256 | > 256 | > 256 | > 256 | > 256 | > 256 | > 256 | > 256 | > 256 |
| B - RC determined at the susceptibility breakpoint of meropenem (2 mg/L) (mg/L) |  |  |  |  |  |  |  |  |  |  |  |
|  | *Acinetobacter baumannii* |  |  | *klebsiella pneumoniae* |  |  |  | *Escherichia coli* |  |  |  |
| Compound | NCTC 12156 (No MBL) | DT05 44 (NDM -1) | DT011 39 (NDM-1) | NCTC134 42 (No MBL) | NTC134 43 (NDM-1) | ArmMLGA 211 (NDM-1) | ESSCAI1 22 (NDM-1) | SM247682 033 (No MBL) | MLGA 91 (NDM-5) | MLGA 92 (NDM-5) | ESCAI2 31 (NDM-5) |
| Pyruvic acid | < 1 | > 256 | > 256 | < 1 | > 256 | > 256 | > 256 | < 1 | > 256 | > 256 | > 256 |
| 3-Bromopyruvate | < 1 | 32 | 32 | < 1 | 128 | 32 | 64 | < 1 | 32 | 32 | 16 |

Control isolates without MBLs were found to be susceptible to treatment with meropenem, with an MIC of < 1 mg/L being observed. In contrast, isolates with MBLs NDM-1 or NDM-5 demonstrated a marked reduction in susceptibility to meropenem. For the meropenem resistant strains, MIC values ranged from 64 mg/L to 128 mg/L, with one isolate exhibiting an MIC > 256 mg/L. When the isolates were treated with pyruvic acid or 3-BP alone, no measurable activity was observed (> 256 mg/L), indicating a lack of intrinsic antimicrobial activity of these two compounds.

3-BP was able to restore the antibacterial activity of meropenem, set at the EUCAST susceptibility breakpoint of 2 mg/L against resistant isolates and reported as the rescue concentration (RC) (**Table 1B)**. In general, a rescue concentration of 32 mg/L 3-BP was required to restore the activity of meropenem against the tested isolates, independent of the presence of NDM-1 or NDM-5. It was previously reported that 3-BP showed inhibition of NDM-1 in cell-based assays, with 3-BP (64 mg/L) restoring the activity of both meropenem and cefazolin against *E. coli* strains producing NDM-1.^21^ Our observations show that 3-BP exhibits enhanced activity against NDM-1 producing *A. baumannii* isolates (DT0544 and DT01139) and NDM-5 producing *E. coli* isolates (MLGA91, MLGA92 and ESCAI231), requiring 32 mg/L 3-BP to restore meropenem activity. *Klebsiella pneumoniae* isolates required the highest concentration of 3-BP to restore meropenem activity, with two out of the three MBL containing isolates requiring more than 32 mg/L 3-BP, with NCTC13443 requiring 128 mg/L and ESSCAI122 requiring 64 mg/L 3-BP to restore meropenem activity. Pyruvic acid, the parent scaffold of 3-BP displayed no significant activity against any of the isolates containing genes encoding for NDM-1 or NDM-5, consistent with reported results against NDM-1.^21^ These observations support the proposal that the B1 MBL inhibitory activity of 3-BP is due to the reactivity of its electrophilic halomethyl ketone.

3-BP was also investigated as a potential SBL inhibitor, with minimum inhibitory concentrations (MICs) of amoxicillin, pyruvic acid and 3-BP determined according to the European Committee on Antimicrobial Susceptibility Testing (EUCAST) guidelines (**Table 2A)**.^28^ Isolates of *Enterobacter cloacae, K. pneumoniae* and *E. coli* producing at least one of following SBLs; TEM-1B, CTX-M-15, ACT-16, OXA-1 or OXA-48 were tested against (**SI-2**).^30^ Additional isolates were obtained through the National Collection of Type Cultures and noted by their NCTC number.

**Table 2.** (**A**) Microbiological comparison of amoxicillin, pyruvic acid and 3-bromopyruvate activities against SBL containing *E. cloacae, K. pneumoniae and E. coli* strains. MIC: minimum concentration required for no visible growth of bacteria. (**B**) Concentration of pyruvic acid and 3-BP needed to restore the activity of amoxicillin at its EUCAST susceptibility breakpoint concentration of 8 mg/L in SBL containing *E. cloacae, K. pneumoniae* and *E. coli* strains. RC: minimum concentration required for no visible growth of bacteria when used synergistically with amoxicillin set at a concentration of 8 mg/L. Plates were incubated at 37°C for 16–18 h. Three biological and three technical replicates were performed. * - Table 2B, value of > 256 mg/L corresponds to no observable recovery of meropenem activity at 2 mg/L.

| A - MIC (mg/L) |  |  |  |  |  |
| --- | --- | --- | --- | --- | --- |
|  | *Enterobacter cloacae* | *klebsiella pneumoniae* | *Escherichia coli* |  |  |
| Compound | SM247691861A<br>(ACT-16) | NCTC13442<br>(TEM-1B, OXA-48, SHV-89) | SM247682033<br>(TEM-1B) | SM247592071<br>(TEM-1B, CTX-M-15) | SM247601654<br>(CTX-M-15, OXA-1) |
| Amoxicillin | 256 | 256 | 256 | 256 | 256 |
| Pyruvic acid | > 256 | > 256 | > 256 | > 256 | > 256 |
| 3-Bromopyruvate | > 256 | > 256 | > 256 | > 256 | > 256 |
| B - RC determined at the susceptibility breakpoint of amoxicillin (8 mg/L) (mg/L) |  |  |  |  |  |
|  | *Enterobacter cloacae* | *klebsiella pneumoniae* | *Escherichia coli* |  |  |
| Compound | SM247691861A<br>(ACT-16) | NCTC13442<br>(TEM-1B, OXA-48, SHV-89) | SM247682033<br>(TEM-1B) | SM247592071<br>(TEM-1B, CTX-M-15) | SM247601654<br>(CTX-M-15, OXA-1) |
| Pyruvic acid | > 256 | > 256 | > 256 | > 256 | > 256 |
| 3-Bromopyruvate | > 256 | > 256 | > 256 | > 256 | > 256 |

Isolates bearing SBLs manifested resistance to amoxicillin, with isolates exhibiting an MIC of 256 mg/L, in contrast to the observed meropenem susceptibility (MIC of < 1 mg/L) of the strains NCTC13442 and SM247682033. When the isolates were treated with pyruvic acid or 3-BP alone, no measurable activity was observed (> 256 mg/L), again indicating a lack of intrinsic antimicrobial activity. Neither pyruvic acid nor 3-BP restored the activity of amoxicillin set at the EUCAST susceptibility breakpoint of 8 mg/L, with MIC values exceeding 256 mg/L (**Table 2B**). These observations suggest that 3-BP inhibitory activity is selective toward MBLs.

### Mass spectrometry studies

To investigate the role of the active site cysteine in the covalent binding of 3-BP to MBLs, initial small molecule mass spectrometry studies were performed against cysteine and glutathione to ascertain if 3-BP readily forms a covalent bond with biologically relevant sulphur containing compounds as previously reported.^31^ As anticipated, mass spectrometry studies demonstrated the reaction of cysteine and glutathione with 3-BP, giving S-alkylated products. Two conjugate mass peaks of 208.0311 Da and 394.0911 Da were obtained respectively (**Figure 1**).

**Figure 1.**
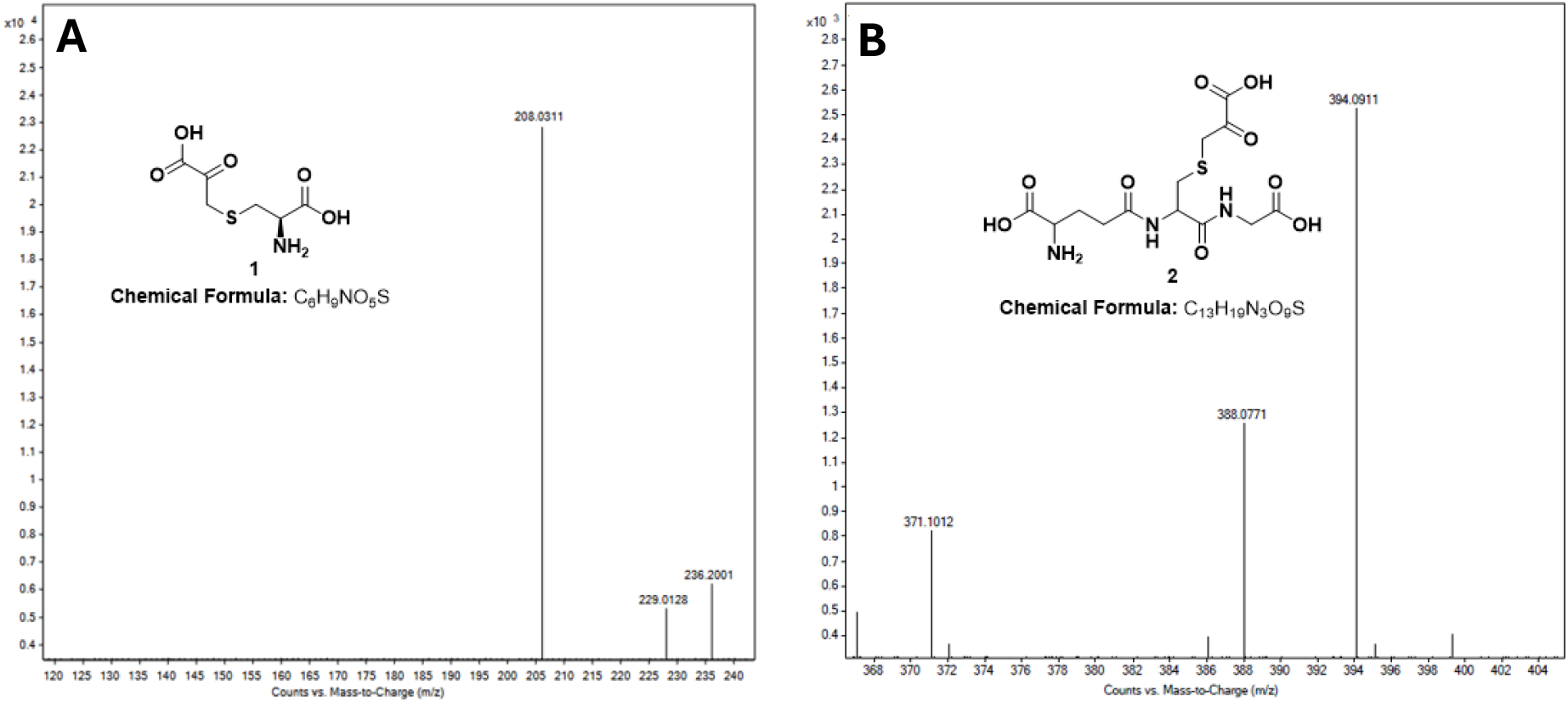
Small molecule mass spectrometry assays show reaction of 3-BP (10mM) conjugation with cysteine (10mM) 208.0311 Da (**A**) and glutathione (10mM) 394.0911 Da (**B**) under conditions used to replicated those of MIC assays (37°C, 16–18 h).

The reaction of 3-BP with Zn(II) complexed- and apo-(non-zinc containing) NDM-1 and NDM-5 was investigated using denaturing electrospray ionisation mass spectrometry conditions. When comparing the amino acid sequences of each MBL, only one cysteine can be found in each enzyme, meaning that if any 3-BP were to bind with a solvent accessible nucleophile, it would bind at the Cys(208) position within the active site and not at an allosteric site. AlphaFold was used for tertiary protein structure modelling of the two apo-proteins.^32^ It was found that both NDM-1 and NDM-5 shared the same B1 MBL subclass active site as expected, with NDM-5 varying from NDM-1 through two amino acid substitutions at position 88 (Val-Leu) and position 154 (Met-Leu) as highlighted in purple (**Figure 2**). These substitutions have resulted in NDM-5 showing an increased resistance and stability under conditions where zinc is depleted.^33, 34^ This was observed during buffer apo-enzyme preparation, where it was noted that NDM-1 precipitated from solution in the absence of zinc, yet NDM-5 remained in solution.

**Figure 2.**
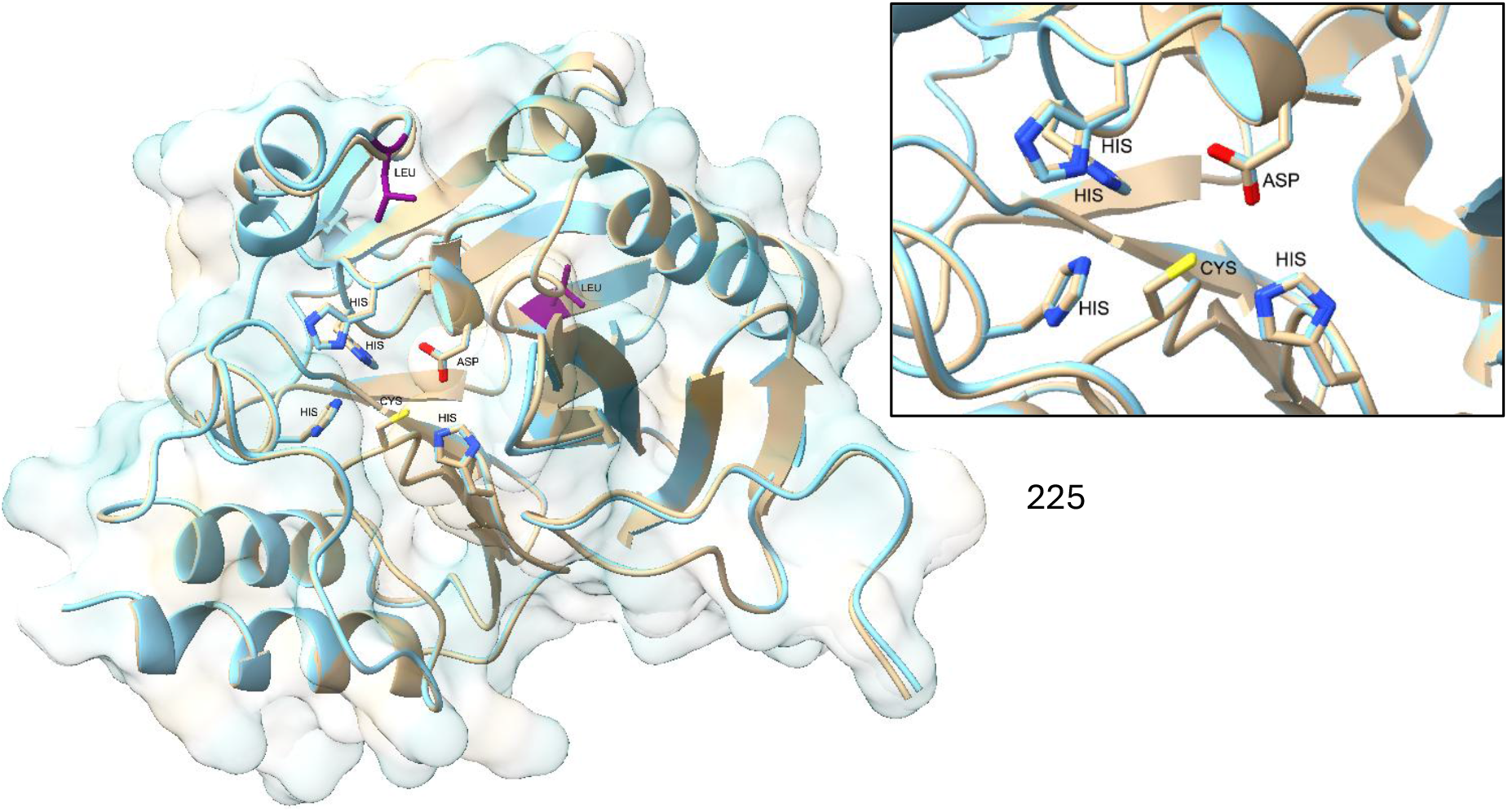
Comparison of AlphaFold modelled apo-NDM1 (brown) and NDM-5 (blue). Overlayed views show high structural similarity with changes in NDM-5 highlighted in purple, (Val88Leu and Met154Leu). **Insert**, expansion of the apo-NDM-1 (brown) and NDM-5 (blue) active sites. The zinc-1 site is defined by three histidine residues whilst the zinc-2 site is defined by a cysteine, histidine and aspartic acid residue.

Ambler class B1 MBLs contain two Zn (II) cofactors bound in the zinc-1 and zinc-2 sites where the latter is reported to show reduced Zn (II) binding affinity compared to that found in the with zinc-1 site.^35-37^ This zinc-2 site contains a conserved cysteine residue (Cys_208_ in NDM-1), therefore, if the zinc-2 site becomes vacant upon dissociation of the Zn(II) metal ion, a free thiol on the cysteine would become exposed and be accessible to modification by 3-BP.^38^ 3-BP was tested against both the natural- and apo-(non-zinc containing) form of purified MBLs, NDM-1 and NDM-5, each of which contain only one cysteine, under denaturing mass spectrometry conditions. 3-BP reacted with both NDM-1 and NDM-5 regardless of whether Zn(II) was present (**Figure 3**).

**Figure 3.**
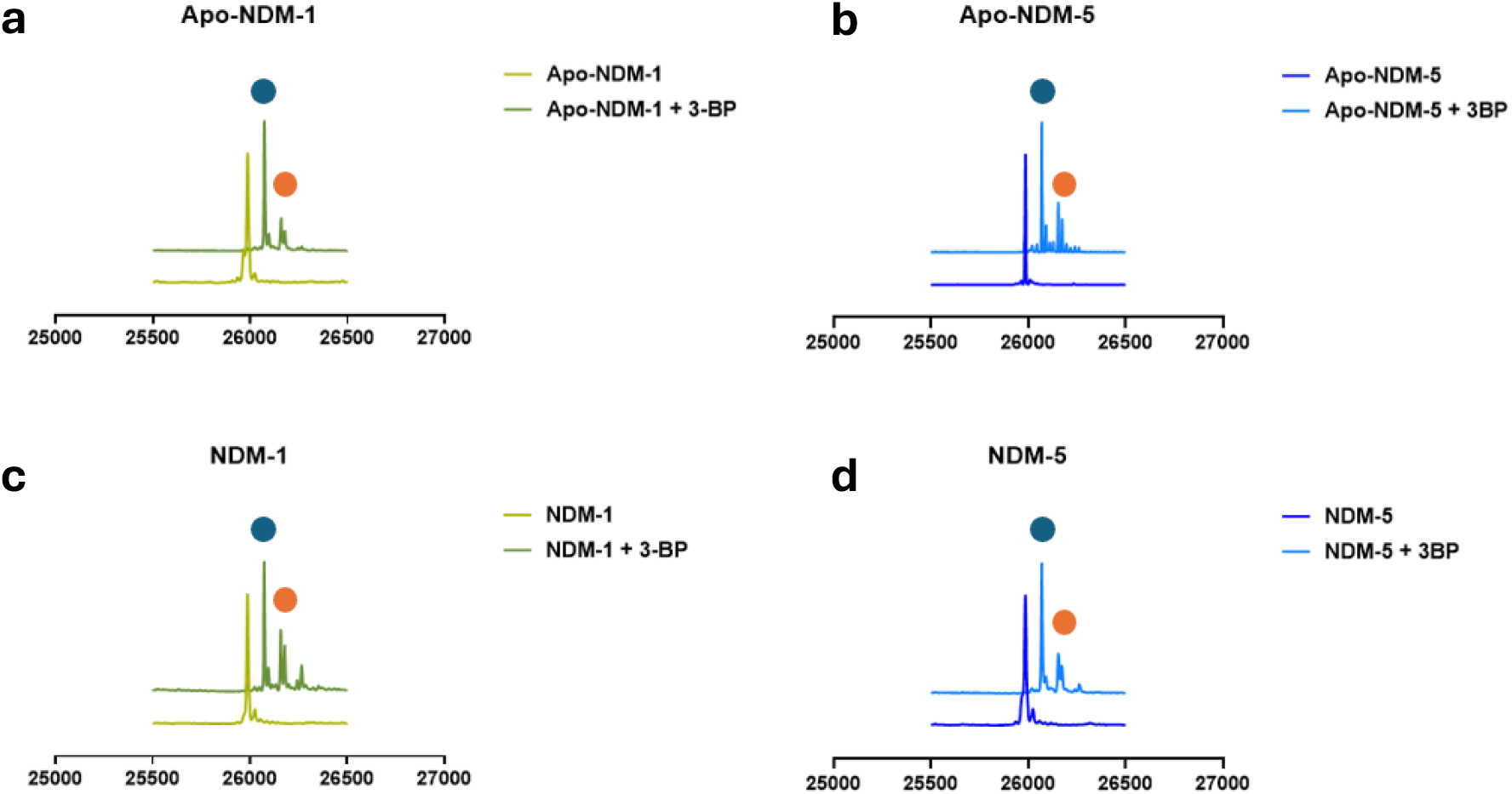
Denaturing mass spectrometry results of apo-NDM-1/NDM-5. (**a, b**) and in the presence of Zn(II) (**c, d**). Control spectra without 3-BP present is shown as the baseline. NDM-1 and NDM-5 have an observed mass of 25987 Da and 25986 Da. A thioether adduct **6** was formed with NDM-1 and NDM-5 in both the apo- and zinc metalled enzymes highlighted by the blue circle (+ 87 Da). The presence of additional peaks (orange circle) is proposed to be an intact tetrahedral hemithioketal with fragments identified as + 166 Da and + 168 Da, proposed to be ^79^Br and ^81^Br isotopes in the intact 3-BP structure **4**. An additional fragment at +171 Da was also observed, potentially also being the intact adduct or the formation of two alkylation events. A fragment at + 281 Da was unable to be identified in NDM-1 and was assigned as an artifact.

The observed mass for NDM-1 and NDM-5 was found to be 25987 Da and 25986 Da respectively and in good agreement with the calculated mass of 25988 Da and 25984 Da. It was noted that as denaturing mass spectrometry conditions were used, only apo-proteins were observed. Upon the addition of 3-BP to both the apo- and metallated NDM-1 and NDM-5, the major fragment observed was a + 87 Da fragment, resulting in a mass (NDM-1 = 26074 Da, NDM-5 = 25986 Da) corresponding to a covalent reaction with 3-BP, resulting in HBr loss and the formation of covalent adduct **6** (**SI-3**). In both enzymes, additional fragments were also observed of + 166 Da and + 168 Da, potentially corresponding to the ^79^Br and ^81^Br isotopes of an intact covalent 3-BP adduct (26153 Da and 26155 Da in NDM-1, 26152 Da and 26154 Da in NDM-5). Although further work is required the additional fragments may correspond to a hemithioketal (**4)**, consistent with an initial reaction with the ketone of 3-BP (**Figure 4**).^39^

**Figure 4.**
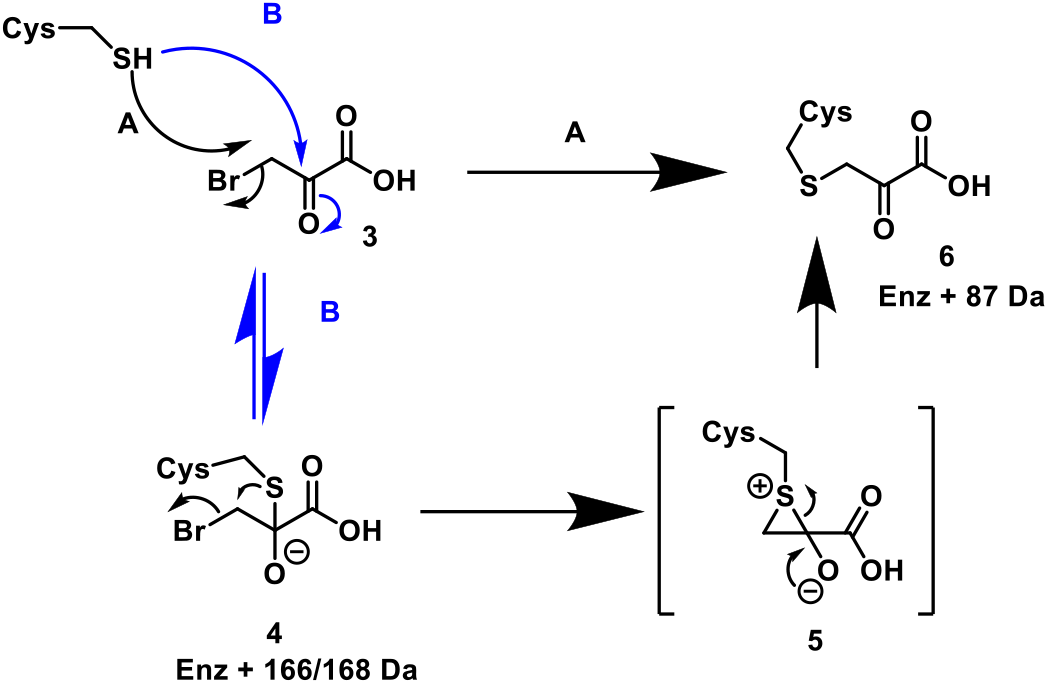
Possible pathways for the reaction of 3-bromopyruvate covalent bond formation with active site cysteine residue. Pathway **A** shows S_N_2 displacement of the bromide giving a thioether. Pathway **B** shows initial reaction at the ketone, followed by rearrangement to give the thioether adduct. Possible mass spectrometry fragments noted their enzyme + conjugate fragment.

Similar observations have been made with the reactions of halomethyl ketones against other nucleophilic cysteine containing proteins, with crystallography data supporting the formation of a hemithioketal intermediate.^39-42^ Previously it was proposed that 3-BP forms a reversible covalent adduct with NDM-1 via pathway **A**; it is, however, generally considered that reactions of halomethyl ketones (such as 3-BP) with cysteines are irreversible, due to the formation of a stable thioether bond.^21, 39, 42^ As a result, it is proposed that pathway **B** and the formation of the intermediate hemithioketal is more likely to reflect an initial reversible covalent reaction previously reported, followed by irreversible rearrangement to give a thioether.^43, 44^

3-Bromopyruvate displayed here shows solid foundation to be built upon, showing evidence that MBLs can be targeted covalently and may allow for the development of future clinically useful MBL inhibitors. In summary, 3-bromopyruvate was shown to inhibit NDM-1 in *A. baumannii* and *K. pneumoniae* and NDM-5 in *E. coli* isolates, expanding on previous findings.^21^ Microbiological studies showed that 3-BP alone did not have any intrinsic activity but restored the activity of meropenem (2 mg/L) against a total of eight resistant MBL producing clinical or environmental isolates of *E. coli, K. pneumoniae and A. baumannii* when co-administered. 3-BP failed to restore the activity of amoxicillin (8 mg/L) against five SBL producing isolates of *E. coli, K. pneumoniae and E. cloacae*, reinforcing the potential of 3-BP as a selective MBL inhibitor. The parent scaffold, pyruvic acid showed no inhibitory activity against MBL or SBL producing isolates, meaning the inhibitory activity of 3-BP against MBLs came from the addition of bromine to the scaffold and the introduction of covalent interactions with the MBL active site cysteine.

MS studies evidence a covalent reaction between 3-BP and an active site cysteine to irreversibly give a thioether, potentially via a reversibly formed hemithioketal. 3-Bromopyruvate (3-BP) is a hexokinase inhibitor and despite its reactivity has been explored as a potential anticancer treatment.^22-24^ Studies with mouse fibroblastic cells indicate 3-BP has low cytotoxicity at up to 200 μM.^21^ Therefore, although its reactivity may preclude its development, it may inspire the development of covalently reacting inhibitors for MBLs, an inhibition strategy that has been successful for SBL inhibitors. Likely reflecting the sometimes-advantageous properties of covalent inhibition, including prolonged target inactivation and, in many cases, irreversible, enabling efficient substrate competition ^45-48^

## Author Statements

### Authors’ Contributions

Conceptualisation JKB, DH & APR. Investigation JKB & KCT. Data curation JKB & KCT. Resources SJM, ES, PN, NAF, APR, DH & CJS. Writing – original draft JKB. writing – review & editing all authors. Supervision APR, NAF, DH, CJS. Funding acquisition APR, DH & CJS.

## Acknowledgements

We thank Nicholas A. Feasey for sharing isolates via the JPIAMR funded STRESST consortium.

## References

1. Prestinaci, F.; Pezzotti, P.; Pantosti, A., Antimicrobial resistance: a global multifaceted phenomenon. Pathog Glob Health 2015, 109 (7), 309–318.

2. Murray, C. J. L.; Ikuta, K. S.; Sharara, F., Global burden of bacterial antimicrobial resistance in 2019: a systematic analysis. The Lancet 2022, 399 (10325), 629–655.

3. Mihelič, M.; Vlahoviček-Kahlina, K.; Renko, M.; Mesnage, S.; Doberšek, A.; Taler-Verčič, A.; Jakas, A.; Turk, D., The mechanism behind the selection of two different cleavage sites in NAG-NAM polymers. IUCrJ 2017, 4 (Pt 2), 185–198.

4. Papp-Wallace, K. M.; Endimiani, A.; Taracila, M. A.; Bonomo, R. A., Carbapenems: past, present, and future. Antimicrob Agents Chemother 2011, 55 (11), 4943–60.

5. Zapun, A.; Contreras-Martel, C.; Vernet, T., Penicillin-binding proteins and β-lactam resistance. FEMS Microbiology Reviews 2008, 32 (2), 361–385.

6. Drawz, S. M.; Bonomo, R. A., Three Decades of β-Lactamase Inhibitors. 2010, 23 (1), 160–201.

7. Bush, K.; Bradford, P. A., β-Lactams and β-Lactamase Inhibitors: An Overview. 2016, 6 (8).

8. Crowder, M. W.; Spencer, J.; Vila, A. J., Metallo-β-lactamases: Novel Weaponry for Antibiotic Resistance in Bacteria. Accounts of Chemical Research 2006, 39 (10), 721–728.

9. Galleni, M.; Lamotte-Brasseur, J.; Rossolini, G. M.; Spencer, J.; Dideberg, O.; Frère, J.-M., Standard Numbering Scheme for Class B & beta-Lactamases. Antimicrobial Agents and Chemotherapy 2001, 45 (3), 660–663.

10. King, A. M.; Reid-Yu, S. A.; Wang, W.; King, D. T.; De Pascale, G.; Strynadka, N. C.; Walsh, T. R.; Coombes, B. K.; Wright, G. D., Aspergillomarasmine A overcomes metallo-β-lactamase antibiotic resistance. Nature 2014, 510 (7506), 503–506.

11. Ju, L.-C.; Cheng, Z.; Fast, W.; Bonomo, R. A.; Crowder, M. W., The Continuing Challenge of Metallo-β-Lactamase Inhibition: Mechanism Matters. Trends in Pharmacological Sciences 2018, 39 (7), 635–647.

12. Nordmann, P.; Naas, T.; Poirel, L., Global spread of Carbapenemase-producing Enterobacteriaceae. Emerg Infect Dis 2011, 17 (10), 1791–8.

13. Yong, D.; Toleman, M. A.; Giske, C. G.; Cho, H. S.; Sundman, K.; Lee, K.; Walsh, T. R., Characterization of a new metallo-beta-lactamase gene, bla(NDM-1), and a novel erythromycin esterase gene carried on a unique genetic structure in Klebsiella pneumoniae sequence type 14 from India. Antimicrob Agents Chemother 2009, 53 (12), 5046–54.

14. Farhat, N.; Khan, A. U., Evolving trends of New Delhi Metallo-betalactamse (NDM) variants: A threat to antimicrobial resistance. Infection, Genetics and Evolution 2020, 86, 104588.

15. Moyo, S. J.; Manyahi, J.; Hubbard, A. T. M.; Byrne, R. L.; Masoud, N. S.; Aboud, S.; Manji, K.; Blomberg, B.; Langeland, N.; Roberts, A. P., Molecular characterisation of the first New Delhi metallo-β-lactamase 1-producing Acinetobacter baumannii from Tanzania. Transactions of The Royal Society of Tropical Medicine and Hygiene 2021, 115 (9), 1080–1085.

16. Green, V. L.; Verma, A.; Owens, R. J.; Phillips, S. E. V.; Carr, S. B., Structure of New Delhi metallo-[beta]-lactamase 1 (NDM-1). Acta Crystallographica Section F 2011, 67 (10), 1160–1164.

17. Stachulski, A. V.; Schofield, C. J., Small but powerful. Nature Reviews Chemistry 2025, 9 (12), 807–808.

18. Grabein, B.; Arhin, F. F.; Daikos, G. L.; Moore, L. S. P.; Balaji, V.; Baillon-Plot, N., Navigating the Current Treatment Landscape of Metallo-β-Lactamase-Producing Gram-Negative Infections: What are the Limitations? Infect Dis Ther 2024, 13 (11), 2423–2447.

19. Mojica, M. F.; Rossi, M.-A.; Vila, A. J.; Bonomo, R. A., The urgent need for metallo-beta-lactamase inhibitors: an unattended global threat. The Lancet Infectious Diseases 2022, 22 (1), e28–e34.

20. Ihrlund, L. S.; Hernlund, E.; Khan, O.; Shoshan, M. C., 3-Bromopyruvate as inhibitor of tumour cell energy metabolism and chemopotentiator of platinum drugs. Mol Oncol 2008, 2 (1), 94–101.

21. Kang, P. W.; Su, J. P.; Sun, L. Y.; Gao, H.; Yang, K. W., 3-Bromopyruvate as a potent covalently reversible inhibitor of New Delhi metallo-β-lactamase-1 (NDM-1). Eur J Pharm Sci 2020, 142, 105161.

22. Darabedian, N.; Chen, T. C.; Molina, H.; Pratt, M. R.; Schönthal, A. H., Bioorthogonal Profiling of a Cancer Cell Proteome Identifies a Large Set of 3-Bromopyruvate Targets beyond Glycolysis. ACS Chemical Biology 2018, 13 (11), 3054–3058.

23. Cardaci, S.; Desideri, E.; Ciriolo, M. R., Targeting aerobic glycolysis: 3-bromopyruvate as a promising anticancer drug. Journal of Bioenergetics and Biomembranes 2012, 44 (1), 17–29.

24. Ko, Y. H.; Verhoeven, H. A.; Lee, M. J.; Corbin, D. J.; Vogl, T. J.; Pedersen, P. L., A translational study “case report” on the small molecule “energy blocker” 3-bromopyruvate (3BP) as a potent anticancer agent: from bench side to bedside. Journal of Bioenergetics and Biomembranes 2012, 44 (1), 163–170.

25. Yun, Y.; Han, S.; Park, Y. S.; Park, H.; Kim, D.; Kim, Y.; Kwon, Y.; Kim, S.; Lee, J. H.; Jeon, J. H.; Lee, S. H.; Kang, L. W., Structural Insights for Core Scaffold and Substrate Specificity of B1, B2, and B3 Metallo-β-Lactamases. Front Microbiol 2021, 12, 752535.

26. Abramson, J.; Adler, J.; Dunger, J.; Evans, R.; Green, T.; Pritzel, A.; Ronneberger, O.; Willmore, L.; Ballard, A. J.; Bambrick, J.; Bodenstein, S. W.; Evans, D. A.; Hung, C.-C.; O’Neill, M.; Reiman, D.; Tunyasuvunakool, K.; Wu, Z.; Žemgulytė, A.; Arvaniti, E.; Beattie, C.; Bertolli, O.; Bridgland, A.; Cherepanov, A.; Congreve, M.; Cowen-Rivers, A. I.; Cowie, A.; Figurnov, M.; Fuchs, F. B.; Gladman, H.; Jain, R.; Khan, Y. A.; Low, C. M. R.; Perlin, K.; Potapenko, A.; Savy, P.; Singh, S.; Stecula, A.; Thillaisundaram, A.; Tong, C.; Yakneen, S.; Zhong, E. D.; Zielinski, M.; Žídek, A.; Bapst, V.; Kohli, P.; Jaderberg, M.; Hassabis, D.; Jumper, J. M., Accurate structure prediction of biomolecular interactions with AlphaFold 3. Nature 2024, 630 (8016), 493–500.

27. Meng, E. C.; Goddard, T. D.; Pettersen, E. F.; Couch, G. S.; Pearson, Z. J.; Morris, J. H.; Ferrin, T. E., UCSF ChimeraX: Tools for structure building and analysis. Protein Science 2023, 32 (11), e4792.

28. Kadeřábková, N.; Mahmood, A. J. S.; Mavridou, D. A. I., Antibiotic susceptibility testing using minimum inhibitory concentration (MIC) assays. npj Antimicrobials and Resistance 2024, 2 (1), 37.

29. Manyahi, J.; Moyo, S. J.; Kibwana, U.; Goodman, R. N.; Allman, E.; Hubbard, A. T. M.; Blomberg, B.; Langeland, N.; Roberts, A. P., First identification of bla (NDM-5) producing Escherichia coli from neonates and a HIV infected adult in Tanzania. J Med Microbiol 2022, 71 (2).

30. Moyo, S. J.; Manyahi, J.; Blomberg, B.; Tellevik, M. G.; Masoud, N. S.; Aboud, S.; Manji, K.; Roberts, A. P.; Hanevik, K.; Mørch, K.; Langeland, N., Bacteraemia, Malaria, and Case Fatality Among Children Hospitalized With Fever in Dar es Salaam, Tanzania. Frontiers in Microbiology 2020, Volume 11 –2020.

31. Sadowska-Bartosz, I.; Szewczyk, R.; Jaremko, L.; Jaremko, M.; Bartosz, G., Anticancer agent 3-bromopyruvic acid forms a conjugate with glutathione. Pharmacological Reports 2016, 68 (2), 502–505.

32. Abramson, J.; Adler, J.; Dunger, J.; Evans, R.; Green, T.; Pritzel, A.; Ronneberger, O.; Willmore, L.; Ballard, A. J.; Bambrick, J.; Bodenstein, S. W.; Evans, D. A.; Hung, C. C.; O’Neill, M.; Reiman, D.; Tunyasuvunakool, K.; Wu, Z.; Žemgulytė, A.; Arvaniti, E.; Beattie, C.; Bertolli, O.; Bridgland, A.; Cherepanov, A.; Congreve, M.; Cowen-Rivers, A. I.; Cowie, A.; Figurnov, M.; Fuchs, F. B.; Gladman, H.; Jain, R.; Khan, Y. A.; Low, C. M. R.; Perlin, K.; Potapenko, A.; Savy, P.; Singh, S.; Stecula, A.; Thillaisundaram, A.; Tong, C.; Yakneen, S.; Zhong, E. D.; Zielinski, M.; Žídek, A.; Bapst, V.; Kohli, P.; Jaderberg, M.; Hassabis, D.; Jumper, J. M., Accurate structure prediction of biomolecular interactions with AlphaFold 3. Nature 2024, 630 (8016), 493–500.

33. Hornsey, M.; Phee, L.; Wareham, D. W., A novel variant, NDM-5, of the New Delhi metallo-β-lactamase in a multidrug-resistant Escherichia coli ST648 isolate recovered from a patient in the United Kingdom. Antimicrob Agents Chemother 2011, 55 (12), 5952–4.

34. Bahr, G.; Vitor-Horen, L.; Bethel, C. R.; Bonomo, R. A.; González, L. J.; Vila, A. J., Clinical Evolution of New Delhi Metallo-β-Lactamase (NDM) Optimizes Resistance under Zn(II) Deprivation. Antimicrob Agents Chemother 2018, 62 (1).

35. Sychantha, D.; Rotondo, C. M.; Tehrani, K. H. M. E.; Martin, N. I.; Wright, G. D., Aspergillomarasmine A inhibits metallo-β-lactamases by selectively sequestering Zn2+. Journal of Biological Chemistry 2021, 297 (2).

36. Rasia, R. M.; Vila, A. J., Exploring the role and the binding affinity of a second zinc equivalent in B. cereus metallo-β-lactamase. Biochemistry 2002, 41 (6), 1853–1860.

37. Murphy, T. A.; Catto, L. E.; Halford, S. E.; Hadfield, A. T.; Minor, W.; Walsh, T. R.; Spencer, J., Crystal structure of Pseudomonas aeruginosa SPM-1 provides insights into variable zinc affinity of metallo-β-lactamases. Journal of Molecular Biology 2006, 357 (3), 890–903.

38. Kim, Y.; Tesar, C.; Mire, J.; Jedrzejczak, R.; Binkowski, A.; Babnigg, G.; Sacchettini, J.; Joachimiak, A., Structure of apo-and monometalated forms of NDM-1—a highly potent carbapenem-hydrolyzing metallo-β-lactamase. PloS one 2011, 6 (9), e24621.

39. Citarella, A.; Micale, N., Peptidyl Fluoromethyl Ketones and Their Applications in Medicinal Chemistry. Molecules 2020, 25 (17), 4031.

40. Drenth, J.; Kalk, K. H.; Swen, H. M., Binding of chloromethyl ketone substrate analogs to crystalline papain. Biochemistry 1976, 15 (17), 3731–3738.

41. Mittl, P. R. E.; Di Marco, S.; Krebs, J. F.; Bai, X.; Karanewsky, D. S.; Priestle, J. P.; Tomaselli, K. J.; Grütter, M. G., Structure of Recombinant Human CPP32 in Complex with the Tetrapeptide Acetyl-Asp-Val-Ala-Asp Fluoromethyl Ketone*. Journal of Biological Chemistry 1997, 272 (10), 6539–6547.

42. Powers, J. C.; Asgian, J. L.; Ekici, Ö. D.; James, K. E., Irreversible Inhibitors of Serine, Cysteine, and Threonine Proteases. Chemical Reviews 2002, 102 (12), 4639–4750.

43. Bacha, U.; Barrila, J.; Gabelli, S. B.; Kiso, Y.; Mario Amzel, L.; Freire, E., Development of Broad-Spectrum Halomethyl Ketone Inhibitors Against Coronavirus Main Protease 3CLpro. Chemical Biology & Drug Design 2008, 72 (1), 34–49.

44. Otto, H.-H.; Schirmeister, T., Cysteine Proteases and Their Inhibitors. Chemical Reviews 1997, 97 (1), 133–172.

45. Sutanto, F.; Konstantinidou, M.; Dömling, A., Covalent inhibitors: a rational approach to drug discovery. RSC Med Chem 2020, 11 (8), 876–884.

46. Singh, J.; Petter, R. C.; Baillie, T. A.; Whitty, A., The resurgence of covalent drugs. 2011, 10 (4), 307–317.

47. Bauer, R. A., Covalent Inhibitors in Drug Discovery: From Accidental Discoveries to Rational Design. 2015, 1, 1–17.

48. Krishnan, S.; Miller, R. M.; Tian, B., Advances in Covalent Inhibitor Design. 2023, 123, 14210–14265.

