## supporting information for "Covalent Inhibition of New Delhi Metallo-β-Lactamases NDM-1 and NDM-5 by 3-Bromopyruvate"

^e^ Malawi-Liverpool-Wellcome Trust Research Programme (MLW), Blantyre, Malawi.

MBL isolate list

Table 1. Selected clinical or environmental isolates containing metallo-β-Lactamases in combination with other beta-lactamase encoding resistance genes as identified. All other resistance genes not currently identified as of writing as project is still ongoing

|  |  | **Resistance genes** | |
| --- | --- | --- | --- |
| **Bacteria ID** | **ID number** | **Metallo Beta-lactamase gene** | **Other Beta-lactamase genes** |
| *A. baumannii* | DT0544 | *bla*NDM-1  (chromosomal) | *blaADC-25, blaCARB-16, blaOXA-259* |
| *A. baumannii* | DT01139 | *bla*NDM-1 | *blaADC-25, blaCARB-16 and blaOXA-51* |
| *K. pneumoniae* | NCTC 13443 | *blaNDM-1* | blaCTX-M-15, blaCMY-4, blaTEM-1A, blaOXA-9, blaOXA-1 |
| *K. pneumoniae* | ArmMLGA211 | *blaNDM-1* | N/A |
| *K. pneumoniae* | ESSCAI122 | *blaNDM-1* | N/A |
| *E. coli* | ESMLGA91 | *blaNDM-5* | N/A |
| *E. coli* | ESMLGA92 | *blaNDM-5* | N/A |
| *E. coli* | ESCAI231 | *blaNDM-5* | N/A |

SBL isolate list

**Table 2.** Selected clinical or environmental isolates containing serine-β-lactamases in combination with other resistance genes as indicated.

|  |  | **Resistance genes** | | | | |
| --- | --- | --- | --- | --- | --- | --- |
| **Bacteria ID** | **ID number** | **Beta lactam** | **Aminoglycosides** | **Quinolones** | **Cotrimoxazole** | **Others** |
| *E. coli* | 24759-2071 | *blaTEM-1B,*  *blaCTX-M-15* | *aadA5, aph(3'')-Ib, aph(6)-Id* | *-* | *Sul2, sul1,*  *dfrA17* | *mdf(A), mph(A)* |
| *E. coli* | 24760-1654 | *blaCTX-M-15, blaOXA-1* | *aac(3)-IIa, aac(6')-Ib-cr, aph(3'')-Ib, aph(6)-Id* | *aac(6')-Ib-cr* | *sul2, dfrA17* | *tet(B), mdf(A),*  *catB3, catB3* |
| *E. coli* | 24768-2033 | *blaTEM-1B* | *aph(3'')-Ib, aph(6)-Id* | *-* | *sul2, dfrA8* | *mdf(A)* |
| *E. cloacea* | 24769-1861A | *blaACT-16 (AMPC)* | *-* | *-* | *-* | *fosA* |
| *K. pneumoniae* | NCTC 13442 | *blaTEM-1B* *blaOXA-48* blaSHV-89, | *aph(3')-Ia* | - | *sul1, dfrA5, dfrA15* | *tet(*C), catA1, *OqxAB, fosA6, ere(A)* |

**Table 3.** Denaturing mass spectrometry results of apo- and metallated NDM-1 (**A**) and NDM-5 (**B**) in the absence and presence of 3-BP. Observed mass for NDM-1 and NDM-5 was found to be 25987 and 25986 as most abundant mass respectively. Upon addition of 3-BP, the most abundant mass was found to be + 87 Da fragment 26074 Da in NDM-1 and 26073 Da in NDM-5, proposed to the thioether adduct **6**. Additional minor fragments were observed at + 166 and 168 Da corresponding to the ^79^Br and ^81^Br isotopes in the intact 3-BP structure **4**. Due to denaturing conditions used, NDM-1 and NDM-5 were found to not contain zinc in observed spectra. Isotopic distribution of proteins are identified by ± Da difference based on abundance.

| **A-Denaturing mass spectrometry results of NDM-1 with 3-BP** | | | |
| --- | --- | --- | --- |
| **Protein** | **Observed**  **Mass (Da)** | **Δ Mass (Da)** | **Interpretation** |
| **Apo-NDM-1** | 25987 ± 1 | 0 | Control |
| **Apo-NDM-1 + 3-BP** | 26074 ± 1 | + 87 | Major fragment |
|  | 26153 ± 2  26155 ± 2 | + 166  + 168 | ^79^Br minor fragment  ^81^Br minor fragment |
| **NDM-1** | 25987 ± 1 | 0 | Control |
| **NDM-1 + 3-BP** | 26074 ± 1 | +87 | Major fragment |
|  | 26153 ± 2  26155 ± 2 | + 166  + 168 | ^79^Br minor fragment  ^81^Br minor fragment |
| **B-Denaturing mass spectrometry results of NDM-5 with 3-BP** | | | |
| **Apo-NDM-5** | 25986 ± 2 | 0 | Control |
| **Apo-NDM-5 + 3-BP** | 26073 ± 2 | +87 | Major fragment |
|  | 26152 ± 2  26154 ± 2 | +166  +168 | ^79^Br minor fragment  ^81^Br minor fragment |
| **NDM-5** | 25986 ± 2 | 0 | Control |
| **NDM-5 + 3-BP** | 26073 ± 2 | +87 | Major fragment |
|  | 26152 ± 3  26154 ± 3 | +166  +168 | ^79^Br minor fragment  ^81^Br minor fragment |

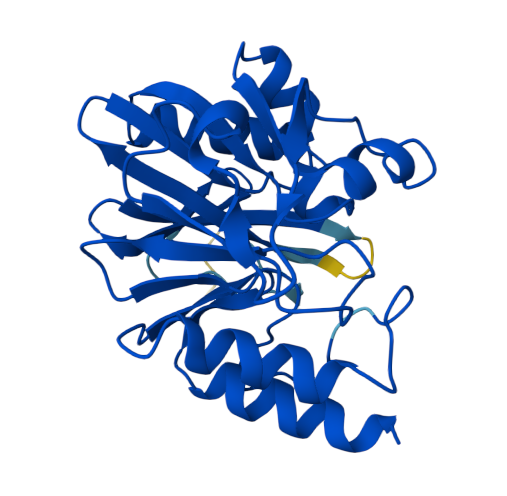

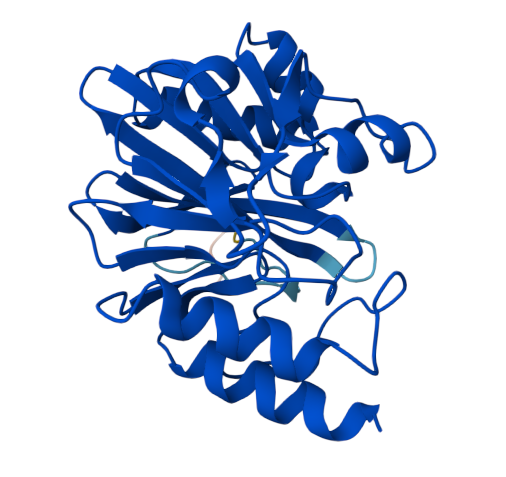

**Figure 1**. AlphaFold model of apo-NDM-1. Coloured by prediction confidence Dark blue (pLDDT > 90), Very high confidence. Light blue (70–90), High confidence. Yellow (50–70), Low confidence. Orange / red (< 50), Very low confidence.

**Figure 2.** AlphaFold model of apo-NDM-5. Coloured by prediction confidence Dark blue (pLDDT > 90), Very high confidence. Light blue (70–90), High confidence. Yellow (50–70), Low confidence. Orange / red (< 50), Very low confidence.

Raw mass spectrometry data available on request.
